# Adaptive learning via surprise gated attractor switching

**DOI:** 10.1101/2025.03.13.643078

**Authors:** Qin He, Daniel N. Scott, Michael J. Frank, Cristian B. Calderon, Matthew R. Nassar

## Abstract

People adjust their use of feedback over time through a process referred to as adaptive learning. We have recently proposed that the underlying mechanisms of adaptive learning are rooted in how the brain organizes time into similarly credited units, which we refer to as latent states. Here we develop a basal ganglia-thalamo-cortical circuit model of this process and show that it captures both the commonalities and heterogeneity in human adaptive learning behavior. Our model learns incrementally through synaptic plasticity in prefrontal-basal ganglia (PFC-BG) connections, but upon observing discordant information, produces thalamocortical reset signals that alter PFC connectivity, driving attractor state transitions that facilitate rapid updating of behavioral policy. We demonstrate that this mechanism can give rise to optimized learning dynamics in the context of either change-points or reversals, and that under reasonable biological assumptions the model is able to generalize efficiently across these conditions, adjusting behavior in a context-appropriate manner. Taken together, our results provide a biologically plausible mechanistic model for adaptive learning that explains existing behavioral data and makes testable predictions about the computational roles of different brain regions in complex learning behaviors.

## Introduction

People and animals exploit the statistics of their environments to optimize behavior through a process that we refer to as adaptive learning (Behrens et al., 2007; Donahue & Lee, 2015). For example, in environments that demand occasionally learning completely new state-action associations, humans and animals prioritize information collected after “changepoints” in the environment to quickly adjust policies corresponding to the new contingency (Li et al., 2019; Nassar et al., 2010). In environments where optimal policy alternates occasionally between two or more alternatives, people and animals can quickly recognize the changes and apply previously learned policies (which we call “reversals” here) (Collins & Koechlin, 2012; Wilson et al., 2014). Bayesian models that reflect knowledge about the structure of the environment have been used to demonstrate the utility of adaptive learning behaviors observed in specific environments (i.e. changepoints and reversals) (Heilbron & Meyniel, 2019; Piray & Daw, 2021; Wilson et al., 2010). However, how adaptive learning is implemented in the brain, and how a fixed neural circuit might implement different forms of adaptive learning, depending on the underlying structure of the environment, remain open and important questions.

One mechanism with potential to achieve adaptive learning across different temporal structures is latent state inference (Collins & Frank, 2013; Gershman & Niv, 2010; Yu et al., 2021). Adaptive learning across a wide range of statistical environments can be cast as a predictive inference problem, in which the solution requires inferring and dynamically updating inferences about the “latent state” – an unobservable variable that gives rise to observable experiences (e.g., task feedback). In this view, partitioning state-action mappings by recruiting distinct neuronal populations for novel states would avoid interference between learning episodes. Moreover, revisiting previously experienced states and reusing established mappings enables faster inference compared to learning from scratch when new experiences closely resemble past ones. While optimal adaptive learning is computationally intractable for complex environments, it can be reasonably well approximated through simplifying assumptions that eliminate the need for attribution of all possible experiences to all possible latent states of the world (Collins & Koechlin, 2012; Fearnhead & Liu, 2007; Lloyd & Leslie, 2013; Nassar et al., 2010; Wilson et al., 2013; Yu & Dayan, 2005).

Neural networks incorporating latent state inference have shed light on a number of behavioral phenomena, even without completely explaining adaptive learning. For example, biologically motivated neural networks have demonstrated that hierarchies of cortico-striatal processing can be used to recruit latent state representations, which in turn constrain downstream action selection (Collins & Frank, 2013; Frank & Badre, 2012). Such models propose how the brain might transfer learning across stimuli and reuse previously learned policies, and explain human behavior across a range of stimulus-response paradigms (Collins & Frank, 2013; Frank & Badre, 2012). In principle, a hierarchy through which prefrontal representations constrain lower-level action selection could also give rise to the full range of adaptive learning phenomena. While this has yet to be tested in a fully elaborated biological network, one recent study used a two-layer feed-forward network to show that representational drift in input layers occurring in response to unexpected feedback at latent state transitions can reproduce normative and human-like adaptive learning behaviors in such tasks (Razmi & Nassar, 2022). This model provided a proof-of-principle, but it abstracted over details that would be critical for implementing the core computations, building them into a real-time biological network, and affording generality across environments (the original model used an “oracle” to dictate how drift should occur in each environment). Thus, there is currently no model that can simultaneously: (i) account for the wide range of adaptive learning behaviors observed in humans, (ii) implement it in a real-time biologically supported circuit, (iii) and generalize across structures (i.e., changepoint/reversals).

To solve these problems, and produce a biologically realistic neural network that accords with human data across tasks, we draw inspiration from empirical observations and well-established computational primitives from systems and computational neuroscience. Each of the building blocks we employ has a long history. Our contribution lies in combining them within a single architecture that affords adaptive learning across temporal structures. First, task-relevant representations are often coded in neural population attractors (Ebitz & Hayden, 2021), which can emerge in recurrent neural networks—classically formalized as auto-associative attractor networks (Hopfield, 1982)—and could potentially be used to store latent state information that persists over time (Brunel, 2003; Hopfield, 1982; Litwin-Kumar & Doiron, 2014; Maes et al., 2020; Recanatesi et al., 2022). Transitions between such attractors can be driven by hetero-associative dynamics (Horn & Usher, 1989), a mechanism we exploit for latent state switching, in our case based on noise (Recanatesi et al., 2022). Second, context information in hierarchically structured tasks is known to be represented within cortico-basal ganglia loops, where corticostriatal plasticity links concrete or abstract states to rewarded actions (Chatham et al., 2014; Collins & Frank, 2013; Frank & Badre, 2012). We build on this framework by implementing plasticity over topographically organized action representations, so that learned PFC-BG mappings guide action selection while action-related activity can also influence PFC representation switches through thalamocortical projections, as shown by experimental studies (Lam et al., 2024; Remington et al., 2018; Wang, Narain, et al., 2018) and supported by theoretical work (Calderon et al., 2022; Recanatesi et al., 2022). Finally, the timing of neural and behavioral state transitions in rodents can be accounted for by dynamic changes in random connectivity that drive activity patterns in recurrent neural networks (Mazzucato, 2022; Recanatesi et al., 2022). These findings collectively imply that attractors storing latent state information in prefrontal regions might be linked to actions through learned associations with the striatum and flexibly recruited by the thalamus altering activity dynamics and facilitating human-like adaptive learning.

Based on the biology just reviewed, we developed a neural model of adaptive learning. Our model can capture human behavior across various tasks and reproduce the emergent behaviors described by Bayesian normative models. Motivated by empirical evidence, it combines active memory representations in prefrontal cortex (Euston et al., 2012; Funahashi & Kubota, 1994; Preston & Eichenbaum, 2013; Rugg et al., 1996; Stokes, 2015; Wall & Messier, 2001) with synaptic plasticity in cortico-striatal connections (Calabresi et al., 2007; Charpier & Deniau, 1997; Fino et al., 2005; Kreitzer & Malenka, 2008; Reynolds & Wickens, 2002). Prefrontal attractors store state representations which become associated with policies through corticostriatal Hebbian learning. Hebbian synaptic plasticity, under this model, is the mechanism implementing supervised learning, which differs from dopamine-based reinforcement learning models of the basal ganglia (while being compatible with them), but is consistent with associative models of corticostriatal plasticity (Charpier & Deniau, 1997; Fino et al., 2005). Supervision itself is a consequence of using the predictive inference tasks we model, which are simpler and more direct measures of switch and update behaviors than reinforcement tasks would be. Given this setting, the model also requires a cortical entry point for instructed information. Primate studies indicate that the frontal eye fields (FEF) and related frontal areas contain both visually responsive and movement-related neurons, making them a natural substrate for integrating sensory feedback with motor plans (Bruce & Goldberg, 1985; Schall, 2002), and our “motor output” layer is meant to represent such frontal populations. Supervision is then processed such that mismatches between expectations and observations trigger thalamocortical transients, which drive prefrontal attractor switches (as seen empirically; Lam et al., 2024; Rikhye, Gilra, and Halassa, 2018; Scott et al., 2024). This model captures a wide range of human behaviors, including both trends and individual differences, and can account for normative aspects of adaptive learning behavior across different conditions and temporal environments. Our results provide insight into how learning and inference, implemented through plasticity and activity dynamics, can be combined in biologically plausible circuits to afford adaptive learning across different temporal contexts.

## Results

Our results detail the development, extension and performance of a biologically plausible model of adaptive learning. We have organized our results as follows. First, we introduce the predictive inference task framework, which has been particularly informative in revealing the computations underlying human adaptive learning. Second, we introduce a model that achieves accurate predictive inference after changepoints by inducing attractor switches and explore how its internal dynamics enable this behavior. Third, we show that a single parameter governing the sensitivity of attractor switches allows the model to capture a wide range of individual differences observed in human behavior. Fourth, we show that adaptively adjusting this parameter through a slight extension of the model enables performance that is robust to noise conditions. Finally, we show that further extending the model to include a feedback projection allows the model to capitalize on repetitive structures without sacrificing performance in non-repetitive task structures.

### Simulated tasks

We tested our model in a predictive inference task environment that has played a primary role in uncovering the dynamics of adaptive learning in humans (Bruckner et al., 2025; Nassar et al., 2010, 2016). The task required participants to predict the next outcome in a series of trials, making behavior minimally complex and maximally informative (McGuire et al., 2014; Nassar & Gold, 2013). In a popular version of the task, a helicopter (occluded by clouds and thus not visible to participants) drops a bag of coins on each trial. Participants are required to catch these bags by positioning a bucket, which serves as a prediction for where the next bag will fall (Figure 1A). After each bag is dropped, participants reposition the bucket according to their updated beliefs about the position of the helicopter, providing an experimental readout of how the most recent bag location affected underlying beliefs about the helicopter location.

**Figure 1.**
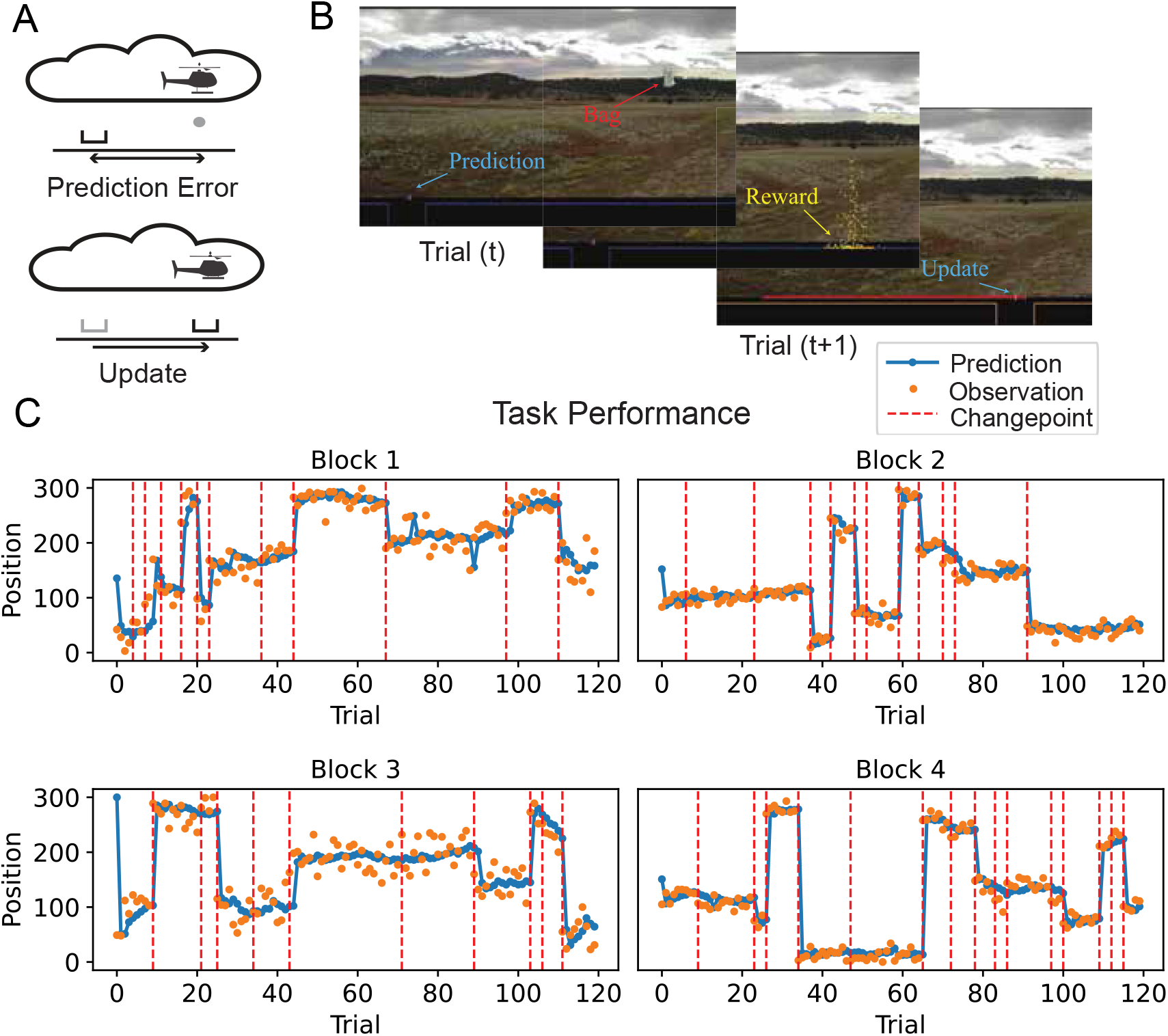
An illustrative example of human behavior in the helicopter task. **A** Cartoon illustration; An occluded helicopter “drops” observations (rewards) and subjects update a “bucket” to catch them, in a trial-wise loop. On each trial, this generates a prediction error and a subsequent update. Participants were incentivized to make good predictions since they earned rewards proportional to the number of coins caught. **B** Annotated screenshots from the task. **C** Predictions from a representative human participant (blue) in response to bag locations (orange) plotted across trials. Bag locations were sampled from a Gaussian distribution, with a mean that was reset at changepoints (red x ticks) and a standard deviation (“noise”) that was experimentally manipulated across blocks of 120 trials. Note that predictions are updated dramatically after changepoints, but much more slowly during periods when the distribution of bags is relatively stable.

In the “changepoint” version of the task, bag locations are sampled from a Gaussian distribution centered on a helicopter location that changes occasionally to an unpredictable new location (Nassar, Bruckner, & Frank, 2019; Nassar et al., 2010). The helicopter location serves as a latent state which must be inferred, and abrupt changes in helicopter location, at so-called changepoints, demand rapid adjustments of behavior. These changepoints render past bag locations irrelevant to the problem of predicting future ones, so that the association of actions with latent states requires that these latent states be partitioned at changepoints. Figure 1B shows an example of the behavior of a human participant performing the helicopter task. Note that the example participant shows the hallmark of adaptive learning in the presence of changepoints, quickly updating predictions after changepoints in the helicopter location but adjusting predictions during periods of stability much more incrementally. Beyond addressing changepoints, predictive inference tasks can be used to reveal adaptive learning behaviors across a much broader range of statistical environments, including reversals and tasks with uninformative latent state noise, thereby providing us with a testbed for generalizing latent state inference mechanisms over environments (D’Acremont & Bossaerts, 2016; Nassar, Bruckner, & Frank, 2019; Piray & Daw, 2024). As we discuss further below, we therefore also run our model on so-called “reversal” tasks, showing that it can also flexibly reuse information in environments containing recurrent contingencies.

### Surprise-induced attractor switch model

Our goal was to build a biologically constrained model capable of achieving human-like adaptive learning in predictive inference tasks. Inspired by empirical observations involving the basal ganglia-thalamocortical circuitry, we propose the **S**urprise-**I**nduced **A**ttractor **S**witch (SIAS) model (Figure 2). The model has a PFC, a BG, and a motor output, with the PFC representing latent states of the environment that the BG transforms into an action selection in the motor layer. Latent states in the PFC are encoded as attractors, the switching of which is controlled by the thalamus. SIAS performs our predictive inference tasks sequentially over trials, by first making decisions based on internal activity, then adjusting internal weights and activity based on feedback. We describe the model elements in more detail using an idealized noise-free changepoint task. Then, we discuss realistic noisy tasks, and the reader is referred to the Methods section for mathematical and implementation details.

**Figure 2.**
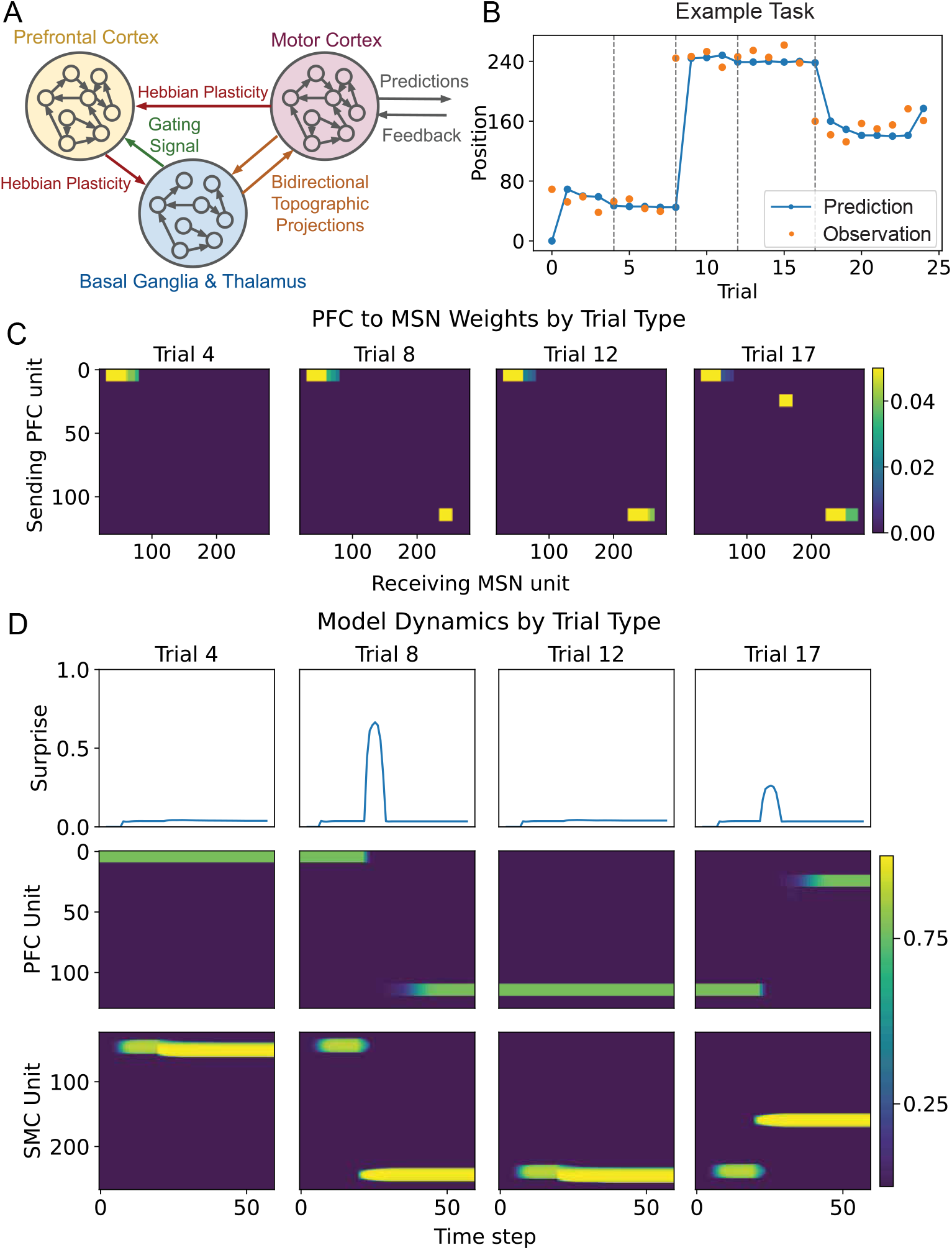
SIAS model architecture and dynamics in a noise-free environment. **A** The SIAS model has three main components. The PFC (yellow) is a Hopfield recurrent network whose attractor states encode latent contexts. It projects to an abstracted basal ganglia-thalamic action-selection circuit (blue), in which topographic Go-like units select actions via learned cortico-striatal associations (red arrow) and shape thalamic activity that relays a surprise-based gating signal back to PFC (green arrow). The motor output layer (purple) is a bump-attractor network that executes actions and receives supervised feedback, which it communicates to the BG and thalamus (orange arrows). In the basic SIAS model, the motor cortex does not project back to the prefrontal cortex. Later, we show that adding this connection enables the flexible model to reuse stored memories for fast inference. **B** Minimal changepoint task illustrating model dynamics. Intra-trial dynamics are shown for the 4 marked (grey vertical line) trials. **C** Evolution of PFC to basal gangalia-thalamus connections over trials. Three attractors at approximate indices 0, 30, and 120 become associated with the three output positions. **D** Surprise signals, PFC activations, and predictions, and switching. Within-trial dynamics evolve over time (x-axis), with predictions occurring at time step 20, followed by a supervised feedback period from timesteps 20–60. Hebbian plasticity is enabled only after an initial delay (timestep 40) within this feedback period, so PFC-BG associations are updated during the later portion of the trial.

The model implements a detailed set of biological circuits, making use of several specific hypotheses about the representations and dynamics in these elements. In the SIAS model, the PFC (yellow area in Figure 2) encodes latent state representations as attractor states within an RNN, which projects to the basal ganglia (BG; blue area in Figure 2) and subsequently to motor outputs (purple area in Figure 2), which collectively perform learning and action selection (Frank & Claus, 2006; Karlsson et al., 2012; Nassar, McGuire, et al., 2019). These attractors are characterized by stable activation of a specific subset of excitatory neurons in the PFC. Attractor states represent specific contexts (i.e., unobservable latent helicopter state). The network then proposes an action associated with each active RNN state via projections to the striatal “Go” cells (Frank & Claus, 2006; O’Reilly & Frank, 2006), which is subsequently relayed through the motor thalamus and ultimately to motor output. We note that our motor output layer is not intended to represent primary motor cortex specifically; rather, it stands for those frontal populations which contain both visually responsive and movement-related neurons, such as the frontal eye fields (Bruce & Goldberg, 1985; Schall, 2002) or adjacent premotor areas, and which could therefore integrate sensory feedback with motor plans. Following action selection, the motor output receives supervised feedback from the environment—encoding the observed bag location in the helicopter task and projects back to the motor thalamus and striatal “Go” cells. This feedback, combined with cortico-striatal plasticity, enables PFC-striatal synapses to learn associations between specific latent states and the observed bag locations.

Model dynamics operate over the sequential unfolding of three main events in each trial: (i) a prediction, (ii) a supervision signal, (iii) a learning window. The core activity and learning dynamics of SIAS can be clearly visualized in a minimal changepoint task that consists of three contexts (or hidden states) with two changepoints (Figure 2B; see below for realistic noisy tasks). To demonstrate this, we analyzed SIAS dynamics in four crucial task transitions, focusing on shifts between stable periods and changepoints (each transition is depicted by a vertical dotted line in Figure 2B). At the behavioral level, the SIAS model quickly adapts to the environment statistics; in particular, the model performs strong updates after changepoints but quickly reduces update magnitudes when responses to consistent latent states are stable, as one would expect from human adaptive learning (Nassar, Bruckner, & Frank, 2019; Nassar, McGuire, et al., 2019; Nassar et al., 2010). Neurally, we observe that in a stable period (e.g., trial 4 or 12) a given attractor state (PFC units in Figure 2D) generates the expected response, without inducing complex state-switching dynamics in SIAS. However, during changepoint events (e.g., trials 8 and 17), SIAS generates an erroneous response, at odds with the subsequent supervision signal. This to-be-resolved (through winner-take-all dynamics) conflict is reflected by an increase in the dissimilarity of thalamus activation (Figure 2D). In turn, this conflict (or surprise signal, see below) elicits a gating signal (green arrow in Figure 2A) that allows PFC state transition by recruiting a novel neuronal population and learning novel state-actions associations.

The way the SIAS model accomplishes the human-like learning just described involves two mechanisms for learning: weight-based plasticity and activation based dynamics (Frank & Claus, 2006; Russin et al., 2025; Wang, Kurth-Nelson, et al., 2018) First, after receiving feedback (motor dynamics, in Figure 2C), cortico-striatal plasticity (red arrow from yellow to blue shaded areas in Figure 2A) allows SIAS to progressively adjust state-action associations. This plasticity follows a Hebbian rule applied to PFC-to-striatum synapses:

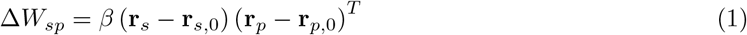

where *β* is the synaptic learning rate, **r**_*p*_ and **r**_*s*_ are PFC and striatal firing rates, and **r**_*p*,0_, **r**_*s*,0_ are pre- and post-synaptic thresholds (see Methods and Table 1). This rule strengthens connections between co-active PFC and striatal neurons, allowing feedback-driven activity to incrementally update the action associated with the currently active PFC representation. We note that these incremental weight adjustments are controlled in magnitude by a synaptic learning rate parameter that is fixed for all trials and simulations - and that this parameter is distinct from our readout of behavioral sensitivity, described in more detail below, which we also refer to as learning rate for historical reasons. Our weights adjustment rule is a simplification of related corticostriatal learning rules in which pre- and post-synaptic activity during the expectation phase can serve as dynamic thresholds, such as the BCM rule (Bienenstock et al., 1982). Whereas reinforcement-learning models typically gate such changes with dopamine reward prediction errors, the predictive inference tasks modeled here use feedback directly changes activity in the motor/BG pathway, updating associations with the currently active, or newly recruited, PFC representation. For simplicity, we use fixed thresholds (Dayan & Abbott, 2005; Sejnowski & Tesauro, 1989), although dynamic activity-dependent thresholds should be compatible with the same architecture.

**Table 1:** Model parameters and default values. A dash indicates the default value is used.

| Symbol | Description | Default | Changepoint | Adaptive | Flexible |
| --- | --- | --- | --- | --- | --- |
| *Network architecture* |  |  |  |  |  |
| $N$ | PFC (RNN) neurons | 600 | — | — | — |
| $P$ | Number of attractor patterns | 60 | — | — | — |
| $N_{\text{patt}}$ | Neurons per pattern ($N/P$) | 10 | — | — | — |
| $M$ | BG / motor output neurons | 300 | 700<sup>†</sup> | — | 700<sup>†</sup> |
| $w_b$ | Bump attractor width (FWHM, neurons) | 15 | — | — | 20<sup>†</sup> |
| $F$ | PFC recurrent weight scaling factor | $P$ | — | — | — |
| $\theta_q$ | Attractor group activation threshold | 0.2 | — | — | — |
| $T_r$ | Rescue delay (steps) | 10 | — | — | — |
| *Layer dynamics* |  |  |  |  |  |
| $\tau$ | Time constant (all RNN layers) | 3.0 | — | — | — |
| $\gamma$ | Self-decay coefficient | 0 | — | — | — |
| $b_s$ | Striatum bias | −0.2 | — | — | — |
| $b_v$ | Thalamus bias | −0.1 | — | — | — |
| $b_o$ | Motor output bias | −0.1 | — | — | — |
| *Inter-layer gains* |  |  |  |  |  |
| $\alpha_{so}$ | Motor output → striatum gain | 5 | — | — | — |
| $\alpha_{vs}$ | Striatum → thalamus gain | 1 | — | — | — |
| $\alpha_{vo}$ | Motor output → thalamus gain | 5 | — | — | — |
| $\alpha_{ov}$ | Thalamus → motor output gain | 1 | — | — | — |
| *Surprise gating* |  |  |  |  |  |
| $S_0$ | Surprise threshold | 0.05 | —<sup>‡</sup> | — | — |
| $I_{\text{inh}}$ | PFC inhibition strength (scalar) | 1.5 | — | — | — |
| *Supervision* |  |  |  |  |  |
| $\eta$ | Supervisory input strength | 5.0 | — | — | — |
| $\sigma_{\text{sup}}$ | Supervisory Gaussian width (FWHM) | 15.0 | — | — | — |
| *Hebbian learning (PFC → striatum)* |  |  |  |  |  |
| $\beta$ | Synaptic learning rate | 0.01 | — | — | 0.02 |
| $\mathbf{r}_{p,0}$ | Pre-synaptic threshold | 0.2 | — | — | — |
| $\mathbf{r}_{s,0}$ | Post-synaptic threshold | 0.2 | — | — | — |
| $W_{\text{min}}$ | Minimum synaptic weight | 0.0 | — | — | — |
| $W_{\text{max}}$ | Maximum synaptic weight | $0.50/N_{\text{patt}}$ | — | — | — |
| $\alpha_d$ | Input competition decay rate | 0.1 | — | — | — |
| $\alpha_s$ | Projection sharpening decay rate | 0.1 | — | — | 0.05<sup>†</sup> |
| $w_s$ | Sharpening half-width (neurons) | 30 | — | — | — |
| *PFC noise* |  |  |  |  |  |
| $\kappa$ | Noise strength | 0.1 | — | — | 1.0 |
| $\tau_{\xi}$ | Noise time constant (steps) | 5.0 | — | — | — |
| $\lambda$ | Noise renewal control | 0.25 | — | — | — |
| *Timing (simulation steps)* |  |  |  |  |  |
| $T_{\text{dec}}$ | Decision phase steps | 20 | 25<sup>†</sup> | — | 25<sup>†</sup> |
| $T_{\text{learn}}$ | Learning phase steps | 40 | — | — | — |
| $T_{\text{delay}}$ | Learning onset delay | 20 | — | — | — |
| *Adaptive threshold (when enabled)* |  |  |  |  |  |
| $\alpha$ | Threshold gain | 2.5 | — | — | — |
| $\tau_{\text{ema}}$ | EMA time constant (trials) | 10 | — | 5 | — |
| *Flexible model (motor → PFC learning)* |  |  |  |  |  |
| $\beta_{po}$ | Learning rate (motor → PFC) | 0.0 | — | — | 0.05 |
| $\alpha_{po}$ | Motor → PFC gain | 1.0 | — | — | 20.0 |
| $\tau_a$ | Active-group decay time constant | 0.0 | — | — | 30.0 |

<sup>†</sup> Figure 6-specific override. Earlier changepoint/adaptive simulations use the default value. <sup>‡</sup> Figure 2 sets $S_0 = 0.1$ for illustration, and Figure 3 sweeps $S_0$ over the fitted threshold range. The Adaptive model uses the changepoint model with adaptive thresholding enabled and exponential EMA updating.

Second, a more pronounced change in learning occurs by altering activity dynamics within the RNN (see second and fourth vertical dashed line in Figure 2B). A surprise signal *S*, computed as a weighted mean absolute deviation of thalamic activity from its spatial centroid, quantifies the agreement between the model’s proposed action and supervised feedback:

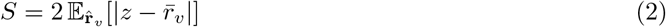

where *z* = *j/M* is the normalized position of neuron *j* over M neurons, 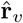 is the L1-normalized thalamic firing rate vector (treated as a distribution over positions), and 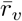 is the activity-weighted mean position. When surprise exceeds a threshold *S*_0_, a gating signal *δ* is activated:

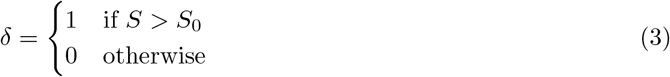

This gating signal controls the external input to the PFC:

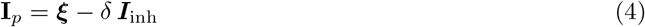

where ***ξ*** is attractor-structured noise and ***I***_inh_ is an inhibition term. When *δ* = 1 (second and fourth columns in Figure 2D), the inhibition suppresses the currently active attractor, allowing noise-driven transitions to a new PFC state—implementing one possible mechanism for dynamics documented in recent experimental work (Halassa & Sherman, 2019; Rikhye, Gilra, & Halassa, 2018; Rikhye, Wimmer, & Halassa, 2018). In particular, that thalamic disinhibition enables noisy inputs to the PFC to facilitate attractor switches. Therefore, weight and activation based dynamics allow SIAS to either integrate new observations into an existing state via Hebbian learning (Eq. (1)), or partition learning into a new PFC latent state and its associated BG connections. Learning can thus proceed incrementally “in-weights” or rapidly via PFC activation dynamics that recruit new PFC populations and cortico-striatal synapses (Frank & Claus, 2006; Russin et al., 2025; Wang, Kurth-Nelson, et al., 2018)

### SIAS mirrors human adaptive learning

People performing predictive inference tasks tend to rely heavily on information immediately following changepoints, but differ in the overall rate of learning, the degree of adjustment at changepoints, and the use of relative uncertainty (which occurs during stable periods). We hypothesized that parameters controlling attractor transitions and Hebbian learning in the SIAS model could recapitulate both trends and differences in human learning behavior. In particular, we expected our threshold parameter for surprise (which may represent balance between stabilizing, inhibitory and destabilizing, disinhibitory thalamocortical inputs (Lam et al., 2024; Mukherjee & Halassa, 2024; Scott et al., 2024)) to account for individual differences in a combination of overall learning and the degree of adjustment to changepoints, while Hebbian learning would impact the degree of learning within context.

Indeed, we found that the model recapitulated group trends in normative learning and that different surprise thresholds in SIAS captured an important source of variation in human learning behavior. Fitting SIAS to subject behavior by adjusting the threshold parameter yielded average adaptive learning dynamics well-matched to those in human participants (Figure 3A). In agreement with normative predictions from Bayesian models, network simulations revealed learning rates, measured as the degree to which predictions are updated for a given prediction error, that were maximal on changepoint trials and decreased as a function of trials thereafter (Figure 3B). However, both the degree of overall learning and its subsequent adjustment depended critically on the surprise threshold, with higher values leading to slower and more stable overall learning rates (Figure 3B; colors refer to different surprise thresholds).

**Figure 3.**
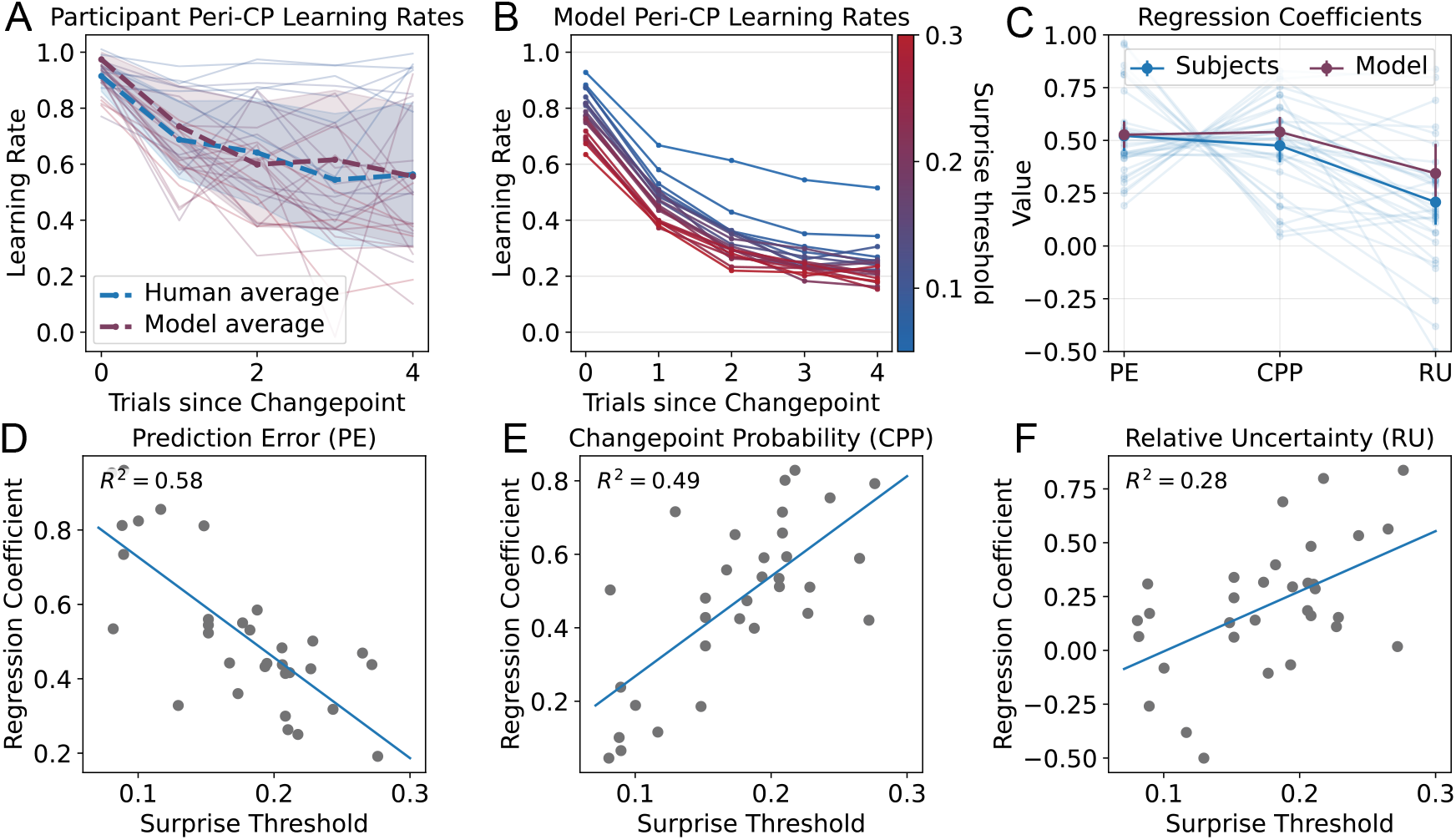
Parameter exploration for surprise threshold and human data regression analysis. **A** We fit individual models (1 free parameter, surprise threshold) to subject data, using MSE on mean perichangepoint learning rates for subjects as objectives. Subject averages and model fits are shown in blue and purple. Each subject’s best-fit result is also shown using the same color scale in panel B. Standard deviations over subjects and models are shown as shaded areas. **B** Applying the model to human data, we ran a parameter sweep over surprise thresholds to illustrate the impacts on learning rate. Learning rate decreases more steeply after changepoints with a higher threshold. **C** Coefficients from a linear regression model, fit on a trialwise basis to both subjects and their best fitting models. Full-opacity blue dots (connected by lines) show average coefficients over subjects. Linear model coefficients fit to best-fit-SIAS-model behavior are shown in purple. Individual subject data are shown in light blue. **D, E, F** Surprise thresholds are correlated with prediction error (**D**), changepoint probability (**E**), and relative-uncertainty (**F**) regression coefficients. The model therefore captures aggregate subject behavior effectively. The individual parameter associations (**D-F**) suggest it captures individual performance characteristics as well, which we also address in the next figure.

Closer examination of model behavior revealed that SIAS adjusted its reliance on recent prediction errors according to normative prescriptions. Normative models of adaptive learning rely on two key factors to control sensitivity to prediction errors (PE): 1) changepoint probability (CPP), accounting for the probability that the helicopter has changed locations since the last observation, and 2) relative uncertainty (RU), accounting for the degree of uncertainty about the current underlying helicopter position. To better understand how SIAS responded to these factors, we fit a linear regression model (see Methods), which predicted updates according to prediction errors along with their interactions with CPP and RU (McGuire et al., 2014; Nassar et al., 2010, 2012; Razmi & Nassar, 2022). SIAS effectively captured the average fixed rate of learning (*β*_*p*_) as well as the degree of learning adjustment according to CPP (*β*_*c*_) and RU (*β*_*r*_) (Figure 3C)(McGuire et al., 2014). Thus, SIAS reproduced normative prediction error-driven learning dynamics, including its modulation by changepoints and subsequent uncertainty, even though prediction errors only enter the model indirectly, through thresholded surprise-gating of PFC attractor transitions.

SIAS not only accounts for overall trends across participants, but also individual differences, including a strong negative relationship between the overall rate of learning and its adjustment according to normative factors (Figure 3C; compare the pattern of gray lines across human and model behavior; when the PE coefficient is high, the CPP and RU coefficients are low, and vice versa). In the case of SIAS, these individual differences are driven by a single factor: the surprise threshold. The PE (*β*_*p*_) coefficient captures the overall learning rate, and participants with higher values of *β*_*p*_ are best fit using lower surprise thresholds (aggressive learner, Figure 3D). CPP (*β*_*c*_) and RU (*β*_*r*_) coefficients are positively correlated with surprise threshold, indicating that participants with higher surprise thresholds (structural learner, Figure 3E-F) more effectively respond to the structure of the task. Therefore, those with higher surprise thresholds make more accurate and conservative use of latent state representations, whereas those with lower thresholds are expected to segregate learning across a greater number of distinct inferred contexts.

Figure 4 gives examples of the SIAS model fit to participants with high and low average learning rates, which are captured using low and high surprise thresholds, respectively. The more aggressive learner makes predictions based almost completely on the most recent observation even during stable periods of the task (Figure 4A,C,E), whereas the more structured learner makes predictions based on previous trials and only adapts to use high learning rates following changepoints (Figure 4B,D,F). For the aggressive learner, learning rates remain consistently at high level regardless of changepoints, whereas for the structured learners, the learning rates start high and gradually decrease to lower levels over time after changepoints (Figure 4E and 4F). SIAS captures behavior on both of these extremes, accounting for not only changes in overall learning rate, but also its dynamics, through adjustments of a single threshold parameter.

**Figure 4.**
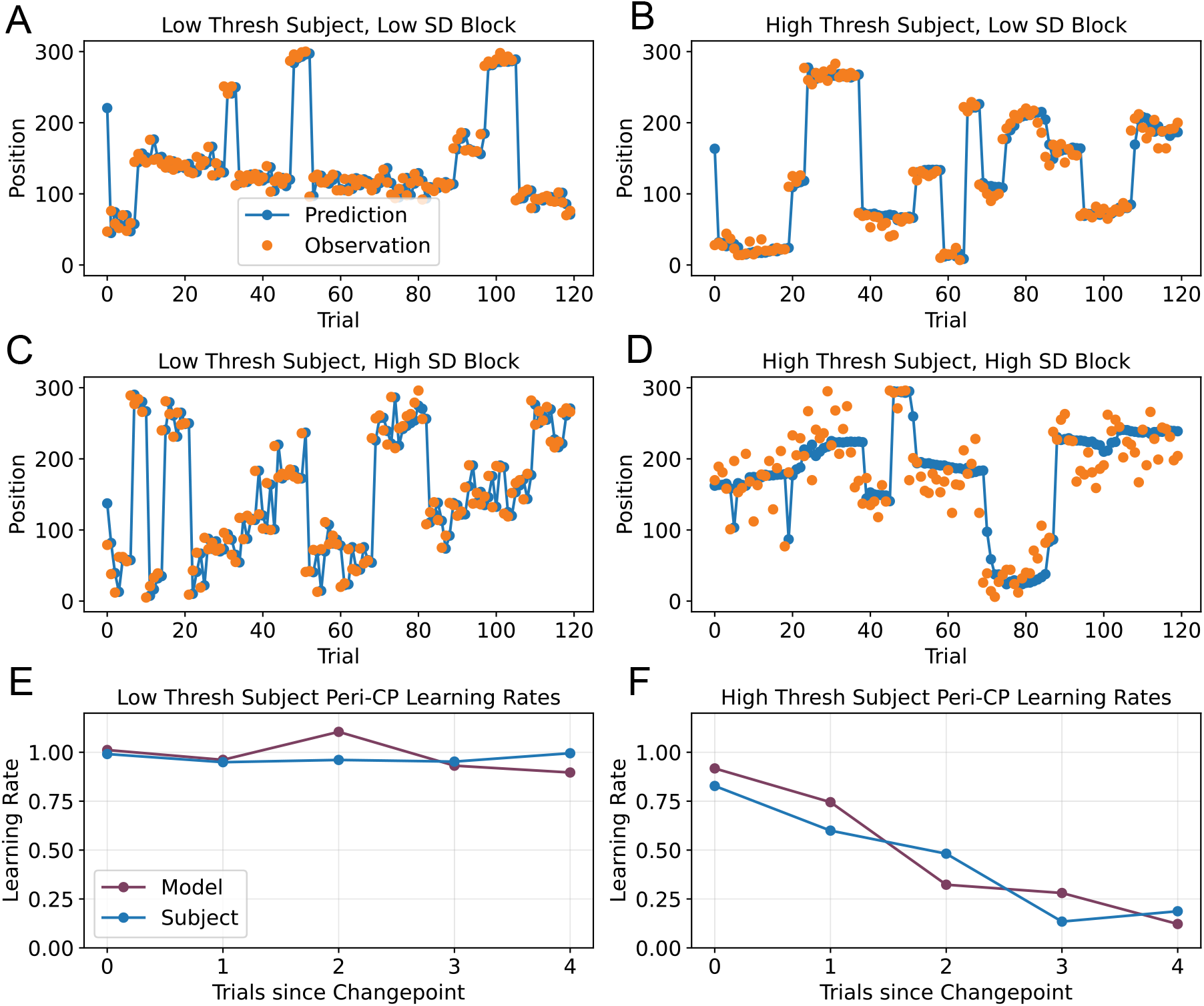
Examples of SIAS models adapt to aggressive learner and structural learner. **A-D** show participant behavior with different learning strategies, respectively. The subject in **A** has a low surprise threshold, and the subject in **B** has a high one. This is especially visible comparing low and high noise blocks (A vs C, B vs D). **E, F** Individual subject model fits against peri-CP learning rates. The model captures both aggregate human learning rate patterns per figure 3, and individual subject behavior, as illustrated here.

While the surprise threshold in the model can be thought of as setting a tolerance for errors beyond which a new latent state is recruited and captures overall learning variability, it imposes a tradeoff between appropriately responding to changepoints and being overly sensitive to noise. As higher surprise threshold reduces noise sensitivity and delays learning in response to changepoints simultaneously. This raises a question of how a single person might adjust their own threshold based on the amount of noise in the current environment(Nassar et al., 2010; Piray & Daw, 2021, 2024), which we address below.

### SIAS captures learning adjustments in response to noise

To mitigate the tradeoff between accurately identifying new states and avoiding excessive sensitivity to noise, we extended the SIAS model to incorporate an adaptive threshold. The fixed threshold simulations above were able to capture individual differences in learning observed in a single noise condition, however, previous work has shown that human participants adjust their learning according to the level of environmental noise. Rather than using a fixed *S*_0_, the adaptive model sets the threshold as a scaled running mean of the surprise observed within each inferred state:

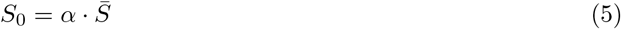

where *α* is a gain parameter and 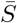 is updated after each trial using an exponential moving average (see Methods and Table 1). This mechanism allows the threshold to rise in high-noise conditions and fall in low-noise ones, adapting the model’s sensitivity to the prevailing level of surprise. By including this adaptive threshold, the model can capture differential learning dynamics across conditions with different levels of noise, as we demonstrate below. Specifically we tested the model in conditions differing in their level of noise and compared the performance of SIAS to that of human participants, as well as to a (reduced, approximate) normative Bayesian inference model (Nassar et al., 2010) with full knowledge of the level of noise in each condition (Figure 5A). This model maintained a belief state at each time based on the computation of changepoint probabilities and outcome expectations conditioned on changepoint occurrence and non-occurrence.

**Figure 5.**
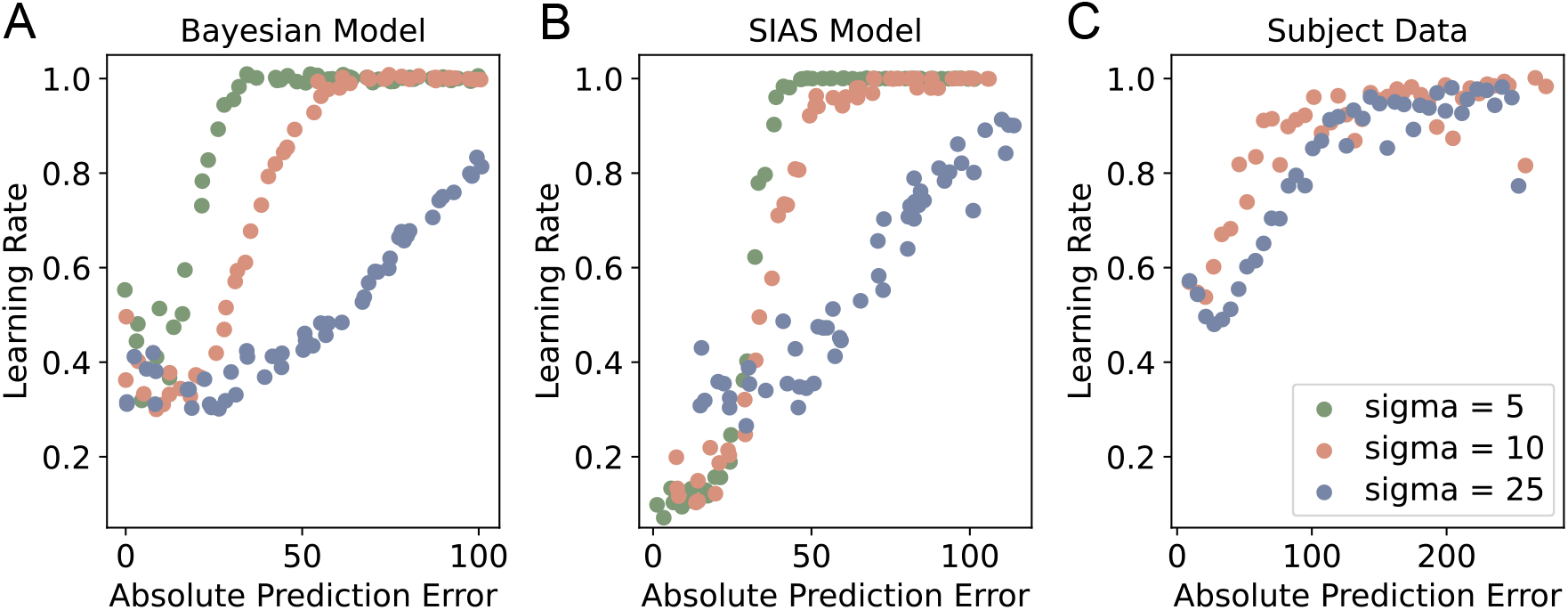
Comparative analysis of models and human performance across varying noise levels. **A** Bayesian ideal observer model. Plots of learning rate by prediction error show that increasing noise flattens the slope of the relationship. **B** SIAS model with adaptive threshold. The adaptive threshold allows the model to adjust its learning rate in response to noise, showing a similar flattening of the learning rate by prediction error relationship as noise increases. **C** Human participants show a similar, albeit attenuated, version of the same pattern. **A-C** All points were computed by binning prediction errors and averaging learning rates within bins. The bins themselves were defined as sliding windows over absolute prediction error. Models were simulated on the same tasks humans performed. All data represent averages over all tasks.

SIAS showed qualitative adjustments to noise prescribed by normative learning to a degree that roughly matched that observed in human participants. Qualitatively, the SIAS model shows a similar dependence on noise as the optimal model in series of generative data (Figure 5B). We pre-trained the model on a sequence of data with a fixed noise level, and then tested on a single prediction error to examine the degree to which it updated in response (defining a learning rate). The adaptive SIAS model required a larger error to increase its learning rate in the face of higher levels of noise, whereas the model without an adaptive threshold presented the same amount of learning behavior for different errors. This adjustment was qualitatively similar to that of a normative learning agent that had full knowledge of the noise conditions, albeit smaller. The adaptive threshold SIAS model shows a shift comparable to that seen in humans across noise levels tested in human experiments (Figure 5C). Taken together, these results suggest that equipping SIAS with an adaptive threshold facilitates normative noise-related adjustments in learning dynamics consistent with human learning.

### SIAS learns new policies and revisits old ones

Humans employ different forms of adaptive learning in different environments. As described above, in situations where changepoints are prevalent, people prioritize experiences at and after such changepoints. However, in environments where contexts are repeated, people often make use of previous learning, for example by recognizing that a previously learned action-outcome contingency would work in a new situation (Collins & Frank, 2013; Yu et al., 2021). This sort of behavior can be measured in tasks that involve contingency reversals, where the appropriate action-outcome mapping occasionally reverts to one that was previously experienced (Collins & Koechlin, 2012; Wilson et al., 2014). The SIAS model described above used surprise-gated functional connectivity to switch during changepoint environments, but had no ability to reuse information after reversals. However, we show in this section that by expanding the scope of learning between areas, SIAS becomes a flexible adaptive learner, learning new policies or reverting to old ones as appropriate given the environmental demands.

We enable SIAS to reuse information, in cases where it would be beneficial, by modifying the machinery governing RNN attractor state transitions. To do so, we allow the RNN attractor state usage to be informed by both new and previously encountered experiences. Specifically, we incorporated feedback projections from the motor output layer directly to the RNN, with weights *W*_*po*_ that are learned slowly through the same Hebbian rule (Eq. (1)). The learned feedback and noise are gated by the surprise signal:

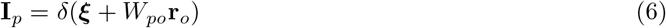

When *δ* = 1, the learned feedback *W*_*po*_ **r**_*o*_ biases transitions toward previously experienced states whose motor outputs match the current feedback, while the noise term continuously drives exploration of novel states. These two mechanisms combine to enable attractor switches that favor novel states when encountering observations unlike those seen before, but previously used ones when encountering observations that closely match previously observed ones.

To assess the flexibility and generality of the augmented model, we compared the previous and the new SIAS models across varying environments. These environments differed in the number of change-points and reversals (Figure 6A). As the ratio of changepoints to reversals increases, the task gradually shifts from having only reversals to mixed scenarios, and ultimately to containing only changepoints. Simulation results from our previous, changepoint-only model show that it was unable to benefit from reversals, achieving similar performance for tasks that were dominated by changepoints and reversals (Figure 6B; purple). However, the flexible version of SIAS that incorporated gated feedback connections was able to achieve higher levels of performance for tasks dominated by reversals, thereby improving on the performance of our original model in these cases (Figure 6B; green). The benefits conferred by our model extensions were related to how the model assigned learning to different PFC attractors. The original SIAS model adjusted weights of a new set of PFC units after each transition, irrespective of whether those transitions reflected changepoints or reversals (Figure 6C; purple). In contrast, the flexible SIAS model scaled the number of PFC attractors used with the number of changepoints, reusing attractors appropriately after reversals (Figure 6C; green). This allows the model to reuse previous knowledge in previously experienced contexts.

**Figure 6.**
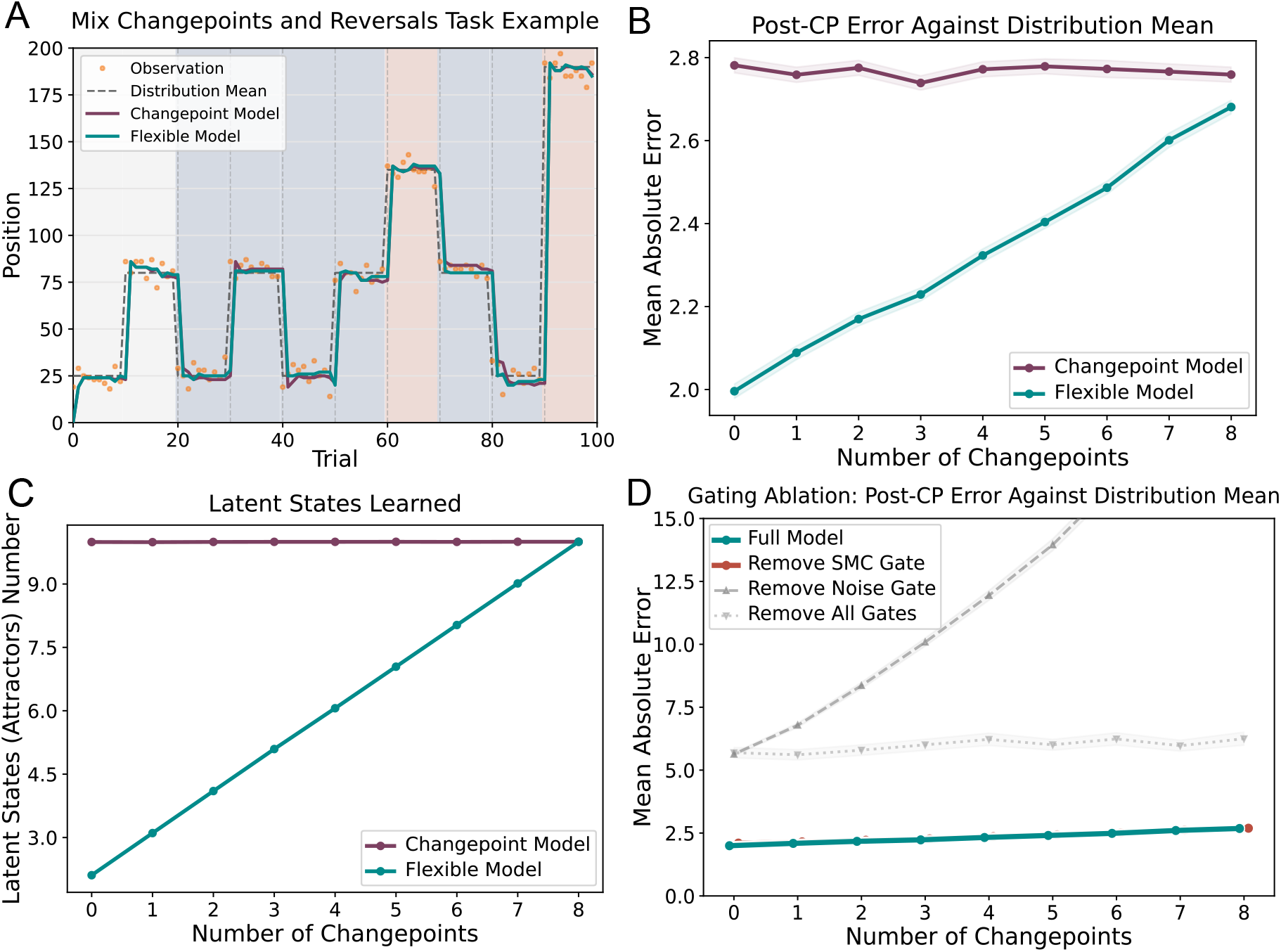
Reversal tasks and flexible model comparison with changepoint model. **A** An example of a predictive inference task including both changepoints and reversals. Outcome position (ordinate) is plotted across trials (abscissa). Each task included two initial warm-up contexts (grey) that were followed by eight subsequent contexts that could include changepoints (red) and/or reversals (blue). Note that this example includes two changepoints and six reversals, but that the full set of tasks included all possible combinations of changepoints and reversals that summed to eight. **B** Mean error following a changepoint or reversal (ordinate) is plotted as a function of the number of changepoints (abscissa), which was inversely proportional to the number of reversals (changepoints + reversals = 8). The flexible model (green) shows performance advantages over the changepoint model (purple) for task environments that are dominated by reversals, rather than changepoints (left). Shading covers ±1 SEM around each mean, with 1000 simulations per point. **C** The improved performance in the flexible model resulted from efficiently mapping learning onto a small number of attractors. The number of attractors that were used (ordinate) is plotted as a function of the number of changepoints (abscissa) for changepoint (purple) and flexible (green) models. Note that the changepoint model uses a new attractor for every switch, irrespective of whether it was a changepoint or reversal, whereas the flexible model reuses previous contexts as appropriate. Points indicate means and shading denotes ±1SEM across simulations but is not visible at this scale. **D** Ablation study of the gating signal reveals importance of noise gates. Mean error following a changepoint or reversal (ordinate) is plotted as a function of the number of changepoints in the task environment for the flexible model (green) and several variants of the model in which surprise gating has been ablated. Note that removing the surprise gate controlling feedback from motor cortex to the PFC had very little impact on model performance (red), whereas removing gating of attractor structured noise (dark gray), or both gates (light gray), had a major negative impact on performance.

To examine the importance of the gating signal in shaping the behavior of the flexible augmented model, we performed an ablation study in which the gate was selectively removed from different components(Figure 6D). In the full flexible model, the gating signal (i.e. surprise) controls both the attractor-structured noise and the cortex -to-PFC connection. Removing the gate on the motor-to-PFC pathway produced performance very similar to the full model, suggesting that this gate is not the main driver of the model’s advantage. By contrast, removing the noise gate substantially impaired performance, indicating that gating the attractor noise is critical for the flexible model. When both gates were removed, noise and motor cortex input were continuously active together, also leading to degradation in model performance, although not quite as severe as in the case where only the noise gate was removed.

Together with the results from the previous sections, our findings show how BG-thalamo-cortical circuitry can afford human-like adaptive learning across environments by combining incremental learning in cortico-striatal synapses with fast inference facilitated by thalamo-cortical projections that drive activity dynamics resulting in selection of context-appropriate cortical populations.

## Discussion

We developed a neural network model, constrained by biology and capable of operating on neuronal timescales, to execute adaptive learning that generalizes across temporal structures. Our SIAS model builds on cognitive models of latent state inference that require two operations: 1) the ability to infer the latent state corresponding to the active action-outcome contingency and 2) the ability to learn appropriate policies for the active latent state (Collins & Frank, 2013; Collins & Koechlin, 2012; Eckstein & Collins, 2020; Gershman & Niv, 2010; Yu et al., 2021). The SIAS model accomplishes these two operations in a corticostriatothalamic architecture whereby inference of the active latent state is accomplished through attractor switches in a prefrontal recurrent neural network and incremental policy learning is accomplished through synaptic plasticity at cortico-striatal synapses. A base version of the SIAS model captures the full range of individual differences in human adaptive learning through adjustment of a single threshold parameter and adjusting this threshold dynamically facilitates human-like adaptive adjustments to environmental noise. Finally, we show that two biologically motivated elements can allow state-revisitation: 1) connections between motor and prefrontal cortex tuned through slow Hebbian learning (Gabbott et al., 2005), and 2) random connectivity in the prefrontal recurrent neural network that is gated by suprathreshold levels of thalamic surprise signaled through thalamo-cortical projections. These elements allow the model to achieve near-optimal inference across environments spanning a range from those that demand reuse of previously learned policies to those that demand rapid relearning in the face of completely new outcome contingencies.

Our model provides a plausible division of labor across cortical and subcortical structures, including cognitive thalamus, that is closely related to several previously investigated ideas. For example, recent work has explored the idea that thalamo-cortical recurrence provides low-dimensional learning pathways complementary to the high-dimensional dynamics of cortex (reviewed in Scott et al., 2024). In computational studies, input from thalamus to cortex has been posited to “switch” cortical dynamics discretely, from implementing one form of processing to implementing another after switch points (Calderon et al., 2022; Hummos et al., 2022, 2024; Lakshminarasimhan et al., 2022; Logiaco et al., 2021). Recordings demonstrating shifts in the computational properties of frontal cortex, mediated by the mediodorsal thalamus (MD), provide empirical support for these ideas (Lam et al., 2025; Mukherjee et al., 2021; Pergola et al., 2018; Scott et al., 2024), and broadly speaking, MD appears to encode context or latent state information, such as task rules or task-sets Mukherjee et al., 2021; Scott et al., 2024). Surprise is a key variable determining when one should attend to one task set versus another, and in line with this observation the frontal thalamus expresses a high density of adrenergic receptors (Pérez-Santos et al., 2021), suggesting that it likely to receive surprise signals at context transitions broadcast from the locus coeruleus (Bouret and Sara, 2005). In our model surprise is used to structure both activity and plasticity, concepts which have been extensively investigated by others (Barry and Gerstner, 2024; Collins and Koechlin, 2012; Liakoni et al., 2021; Scott and Frank, 2024). Unlike dopamine-based reward prediction error accounts that provide a scalar signal of better- or worse-than-expected outcomes, in our model, surprise-based switching is instantiated in surprise-gated attractor transitions that determine when and how upcoming plasticity will be modified. This extends PE-like learning mechanisms to environments where feedback specifies the direction of correction, rather than only the presence or value of an error. The cortico-striatal plasticity in SIAS uses a covariance learning rule, a well-characterized family of Hebbian rules in which synaptic modification depends on deviations of pre- and post-synaptic activity from their respective thresholds (Dayan & Abbott, 2005; Sejnowski & Tesauro, 1989). This formulation naturally produces bidirectional weight changes and selectivity for correlated activity, properties important for SIAS’s ability to form and maintain distinct state-action associations.

Our work also relates to other models proposed to achieve adaptive learning from the perspective of approximate Bayesian inference, and in some cases which have been implemented in BG cortical circuits. Yet previous proposals present several limitations. First, some models require specific assumptions about the environment, such as known generative model parameters, and have not been elaborated in full biological detail (Daw & Courville, 2008; Lloyd & Leslie, 2013; Nassar et al., 2010; Wilson et al., 2013; Yu & Dayan, 2005). As such, they raise questions about both their generality as accounts of adaptive learning and the extent to which they make testable neural predictions. Second, while other models have proposed mechanisms that can either recruit new neuronal populations or reuse previous ones (to either avoid interference or promote knowledge transfer), they have often been tested in deterministic settings that do not incorporate probabilistic features of real-life scenarios or make quantitative predictions for the adaptive learning behaviors examined here (Collins & Frank, 2013; Frank & Badre, 2012). Third, other biologically inspired models have demonstrated the ability to learn adaptively in stochastic environments by adjusting learning rates based on uncertainty, but lack latent state inference, and thus are unable to reuse information in repeating contexts (Franklin & Frank, 2015; Stalnaker et al., 2016). Finally, in an attempt to get closer to neural dynamics, recent work used a two-layer feed-forward network to build state-action pairs with distinct neurons to model human adaptive learning (Razmi & Nassar, 2022). While this model provided a proof-of-principle, it abstracted over details that would be critical for implementing the core computations in real time, and required a structural “oracle” to control how population activity representing state transition structure evolved across trials, and also did not consider repeating contexts. Here, we addressed many of these shortfalls, by presenting a biologically plausible model operating in real time to perform adaptive learning in environments governed by latent states.

An important scope consideration is that the predictive inference tasks we model provide supervised feedback—the true outcome location is revealed on every trial—and SIAS exploits this by routing the supervised signal through the motor output layer to drive both action correction and Hebbian learning in cortico-striatal synapses. This places our model in the supervised learning tradition rather than the reinforcement learning framework more commonly associated with basal ganglia function, in which dopaminergic reward prediction errors modulate striatal plasticity (Frank & Claus, 2006; Schultz et al., 1997). Nevertheless, the core architecture of SIAS – attractor-based state inference in PFC, surprise-gated transitions via the thalamus, and associative learning in cortico-striatal projections – is not inherently restricted to supervised settings. Replacing the supervised motor input with a dopaminergic teaching signal that modulates the same Hebbian plasticity would yield an RL-compatible variant, albeit one that would likely require additional mechanisms for action exploration and temporal credit assignment. We view the supervised case as a natural starting point because it isolates the contributions of latent state inference and associative learning from the additional complexities of reward-based credit assignment, and because the predictive inference tasks that have been the primary empirical testbed for adaptive learning theories provide exactly this type of feedback. As noted above, this also committed us to the establishing an entry point for supervised feedback, which we take to be found in motor areas such as the FEF.

Our model has several implications related to adaptive learning behavior. First, it achieves many of these behaviors, which have traditionally been considered to potentially rely on synaptic learning rate adjustments (Behrens et al., 2007; Browning et al., 2015; Nassar et al., 2010, 2012), without changing synaptic learning rates. Dynamics in the learning behavior of the SIAS model are driven by transitions in a recurrent neural network that provides input to a cortico-striatal learning system. Such transitions allow SIAS to reset policies when entering a new context or to reuse an old policy when entering a familiar one (see Figure 6A). This distinction is important because recent work has highlighted that neural signatures of adaptive learning previously thought to reflect dynamic learning rates (Behrens et al., 2007; Fischer et al., 2015; Nassar et al., 2012) have relationships to behavior that are highly context dependent (D’Acremont & Bossaerts, 2016; Nassar, Bruckner, & Frank, 2019; O’reilly, 2013). Such context dependence is not easy to reconcile with the idea of learning rate adjustment, but is instead better explained by the idea that such signals reflect subjective assessments of latent state transitions (Razmi & Nassar, 2022; Yu et al., 2021). Here we show how such signals, used by SIAS to update prefrontal attractor states, can be harnessed to achieve a wide range of adaptive learning behaviors including some interpretable through the lens of learning rate adjustment (See Figure 3) as well as some that cannot (See Figure 6B).

A second behavioral implication of SIAS is that PFC-BG-Thalamic networks using the well-attested representations and learning rules incorporated in our model should be able to learn efficiently in environments with more complex latent state structure than that assessed here. Latent states learning, when implemented through probabilistic reasoning, requires making assumptions about the underlying structure of the environment, and prior implementations of latent state learning in cognitive and network models have “baked in” the transition structure of the environment in order to achieve behaviors that are efficient for a given task environment. However, people are capable of behaving appropriately across a wide range of different environments, including ones that require rapidly relearning after environmental changes, as well as those that require re-using information learned from an experience in the distant past (Collins & Koechlin, 2012; Nassar et al., 2021). Here we provide a first clue to how people might achieve such flexibility, in that we show how a specific architecture making use of learned feedback projections and gated random connectivity, can support latent state transitions that are appropriate for the current environment, without explicit learning of the environmental transition structure. Here we focus on one specific dimension of transition structure, whether changes in the environment reflect new contexts or repetitions of old ones, and we hope that our work inspires future studies to characterize the breadth of transition structures over which humans can learn, as well as the broader characteristics of network architectures that affect whether and how networks manage learning when confronted with them.

Beyond behavior, SIAS makes specific predictions about its biological basis. We consider some of the larger scale, process-level predictions in this paragraph, and discuss finer ones in the next. A core element of the model is a recurrent neural network that maintains active latent state representations and sends projections to the striatum. A key signature of this sort of representation is that it should change rapidly at transitions in the environment, thereby enabling new subpopulations of neurons to control behavior after these transitions, which minimizes interference between different contexts. While we broadly hypothesize that the attractor network exists in frontal cortex, there is evidence for latent-state like representations in a number of different frontal regions, including dorsolateral prefrontal cortex (Vaidya & Badre, 2022; Vaidya et al., 2021), orbitofrontal cortex (Nassar, McGuire, et al., 2019; Schuck et al., 2016; Wilson et al., 2014), and anterior cingulate cortex (Karlsson et al., 2012; Powell & Redish, 2016). It is noteworthy that all of these regions project to the striatum, albeit to different subregions thereof. Furthermore, “network reset” phenomena subsequent to environmental transitions and tightly linked to behavioral change have been observed in both orbitofrontal cortex in humans (Muller et al., 2019; Nassar, McGuire, et al., 2019) and medial frontal regions of rodents thought to correspond to ACC in primates and humans (Karlsson et al., 2012; Powell & Redish, 2016). Our model proposes that these changes in the cortical representations are provided to the striatum and play a causal role in increasing the rate of behavioral adjustment at environmental changes. This prediction is distinct from other theories based on learning rate adjustment, which suggest instead that periods of rapid learning are accomplished through enhancement of dopaminergic signaling in the striatum (Diederen et al., 2016, 2017; Haarsma et al., 2021), and is broadly in line with the view that an integrated hierarchy of activity and plasticity mechanisms across scales generates metaplasticity that improves behavior (Scott and Frank, 2023). Recent findings that rodent adaptive learning behaviors are not accompanied by scaling of dopamine learning signals (Mah et al., 2024), along with our alternative model of how such adaptive learning behaviors might emerge, suggest the need for new methods that would allow perturbations at the population level to test the causal link between reset of cortical networks and their downstream consequences for learning. Such tests might allow for better distinction between the different cortical regions that have been shown to represent latent states in different tasks and conditions, and potentially reveal their differential contributions to behavior.

The process-level predictions above also yield more specific, experimentally testable claims, which we organize here into functional, experimental, and behavioral categories. To motivate these, recall that SIAS synthesizes four empirically supported observations: (1) frontal recurrent attractors store task representations, (2) cortico-subcortical loops link state and action representations through Hebbian learning, (3) thalamocortical projections switch recurrent states in cortex, and (4) cortical state switching, viewed locally, resembles noise-driven dynamics through structured connectivity. By combining these elements in a single model, SIAS generates functional predictions about how the system is used. Specifically, the model predicts that (A) the task representations maintained in cortical attractors serve as abstract latent variables—contexts—which (B) scaffold additional associative learning by providing a stable basis for cortico-striatal plasticity, while (C) thalamocortical projections use a push-pull mechanism to shut down no-longer-relevant context representations, based on (D) surprise signals computed in the sensory and motor networks linked to action production.

These functional predictions cash out experimentally. First, we expect that different subjects exhibit different surprise thresholds (see Figure 3), and that these thresholds should be directly identifiable as neurometric functions in thalamocortical projection activity—specifically, a sigmoidal relationship between the magnitude of environmental change and the probability of observing a gating signal in frontal thalamic output neurons—and as sigmoidal population similarity curves in PFC, where representational states remain stable for sub-threshold changes and shift abruptly once the threshold is exceeded. Next, the model predicts that associative plasticity between motor-sensory integration populations and recurrent PFC attractors provides the mechanism by which reversal or contextual return happens: Hebbian learning at motor-to-PFC synapses establishes associative links between prior motor output patterns and the PFC attractor states with which they co-occurred, so that familiar sensorimotor activity can bias the PFC toward reinstating the appropriate context. Crucially, this contextual re-activation also requires trans-thalamic surprise signalling to destabilize the currently active attractor, and this requirement is approximately conserved between novel context creation at changepoints and context revisitation at reversals—in both cases the same gating signal initiates the transition, with only the endpoint differing depending on whether learned feedback biases the system toward an existing attractor or noise drives exploration of a new one. Finally, at the behavioral level, eliminating thalamocortical signalling should disrupt context switching while leaving within-context incremental learning intact, degrading performance specifically at context transitions; this deficit should be most pronounced for reversals, where successful performance depends on both the gating signal and the learned motor-to-PFC associations that guide the system back to the correct attractor.

While our model provides a substantial advance in understanding how adaptive learning could be implemented in cortico-striato-thalamic circuitry, it has a number of limitations. First, while our extended architecture is capable of generalization across environments, this generalization is limited to environments with persisting state transitions, such as changepoints and reversals. Thus, while in principle latent state inference could be used to improve learning in environments with sequences or transient outliers (Bakst & McGuire, 2023), accomplishing this in our model would require additional mechanisms for explicit learning of the environmental structure, in particular the transition function defining how latent states are likely to transition to one another. In principle this might be accomplished by extending SIAS through the introduction of plasticity into the RNN, such that gated attractor transitions are biased not only by learned associations to motor outputs, but also through learned associations between latent states that have occurred sequentially in the past.

A second limitation of our model is that it incorporates simplified RNN dynamics and action policies. These simplifications emerged because the goal of our modeling efforts were to explain behavioral data from predictive inference tasks where optimal policies are somewhat simple (report the mean of the underlying distribution) and conditional only on the active generative mean in the absence of trial-specific cues. Here we have attempted to illustrate exactly how combining RNN transitions with a cortico-striatal learning system can be used to facilitate adaptive learning under these restrictive conditions, however the principles that we demonstrate here could be applied to more complex problems such as those faced in other tasks or in real world learning problems. For example, cue information could also be provided to the RNN to yield more complex state representation including both stimulus information as well as its temporal context. Similarly, more complex dynamics could be incorporated into the RNN to yield policies that change over time, or change according to actions initiated. Thus, while we have focused our exploration of the SIAS model on tasks that have been the subject of extensive investigation of behavior and its underlying neural mechanisms (McGuire et al., 2014; Nassar, Bruckner, & Frank, 2019; Nassar, McGuire, et al., 2019; Nassar & Troiani, 2021; Nassar et al., 2010, 2012, 2021), the basic architecture is ripe for extension.

A related open question is how the architecture itself could arise from more fundamental principles, rather than being hand-crafted. Our incremental model progression may offer a clue: the most basic adaptive skill is changepoint detection, which requires only surprise-gated attractor switching and could plausibly emerge from prediction error minimization in recurrent circuits. Context reuse, which requires additional learned feedback, represents a refinement that would confer further advantage only once changepoint detection is already in place. This suggests that the developmental or evolutionary assembly of such a system could recapitulate the incremental structure we have described, with simpler mechanisms providing the scaffold on which more elaborate ones are built.

In summary, the SIAS model provides a biologically motivated account of adaptive learning through latent state inference. It explains human behavior, recapitulates normative adjustments in learning, and can rapidly re-learn or reuse information as appropriate for a given situation. These findings provide a principled explanation for how cortical activity dynamics and striatal plasticity mechanisms can be combined to produce flexible and efficient learning. Future work should test the key biological predictions of our model and build on our results to address more complex statistical environments, and to integrate our model’s account of plasticity with the growing body of related work which suggests that a hierarchy of different mechanisms, distributed within and between areas, supports behavioral plasticity.

## Methods

The SIAS model consists of a PFC that encodes latent states as attractor states within an RNN, projecting to the BG through topographically organized pathways, as implemented in many previous cortico-BG models (e.g., Frank and Claus, 2006; Gurney et al., 2001) and motivated by neuroanatomical data (Alexander et al., 1986). After an action is selected and executed, supervised feedback enters the network, reinforcing existing state-action associations or establishing new ones through plasticity at the PFC-striatal synapses.

In the sections which follow, we first describe the model’s layer dynamics, plasticity rules, and surprise-gating mechanism, including the adaptive threshold and flexible model extensions. We then describe the helicopter task and the analyses used to compare model and human behavior, followed by the mixed changepoint-reversal environments used to evaluate the flexible model. The code and parameter values for all simulations reported here can be found at xxxx (all code will be publicly available after acceptance). Table 1 lists all model parameters and their values.

### The SIAS model

The SIAS model consists of four recurrent neural network layers arranged in a cortico-basal ganglia loop (Figure 2): the PFC (*p*), striatum (*s*), thalamus (*v*), and motor output (*o*). All four layers share the same dynamical equation, differing only in their external inputs:

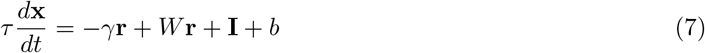

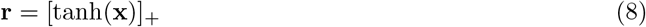

where **x** is the membrane potential vector, **r** is the firing rate vector, *W* is the recurrent weight matrix, **I** is the layer-specific external input, *γ* is a self-decay coefficient, *b* is a bias, and *τ* is the time constant. The rectified hyperbolic tangent guarantees non-negative firing rates throughout. The subsections below specify the external input **I** and recurrent connectivity *W* for each layer; all parameter values are listed in Table 1.

#### PFC dynamics and latent states

The PFC has *N* neurons whose recurrent connectivity implements WTA Hopfield attractor dynamics. The recurrent weight matrix encodes *P* patterns:

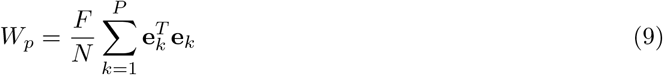

where **e**_*k*_ ∈ ℝ^1*×N*^ is a pattern encoding vector defined as a subset of *N*_patt_ excitatory neurons surrounded by *N* − *N*_patt_ inhibitory neurons, and *F/N* is a scaling factor. The resulting matrix *W*_*p*_ has blocks of excitatory diagonal entries surrounded by inhibitory off-diagonal entries.

At each time step, the mean activation of each attractor group *g* is computed as 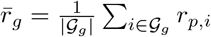 where *G*_*g*_ is the set of neurons belonging to group *g*, and a group is classified as active when the mean activation is above the activation threshold 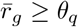. Then, we implemented two mechanisms, quiescence destabilization and co-activation tie-breaking, to ensure that at least one attractor group remains active while destabilizing multiple active groups. Quiescence destabilization is achieved using targeted noise injection into the most-driven group when no group is active for *T*_*r*_ consecutive steps. Co-activation tie-breaking is achieved via destabilizing noise injection when multiple groups are active for *T*_*r*_ steps.

Endogenous noise ***ξ*** is generated as an autocorrelated derivation of a per-attractor point process. Specifically, at each simulation time step, a discrete attractor-aligned noise process samples a candidate attractor group uniformly with *λ* controlling renewal of the noise process. A decay timescale *τ*_*ξ*_ then correlates activity samples over time via exponential convolution. This produces a spatially structured signal aligned with the attractor layout in *W*_*p*_, with intuitive semantics: random attractors push to become spontaneously active at a given rate. For the purpose of exposition, this random noise is represented in the dynamical equation as part of the external input, but it is generated within the PFC layer and thus could be equivalently represented as an additional term in the recurrent component of the dynamics. In changepoint-model simulations, once a PFC attractor became active, future exogenous noise to that attractor was suppressed so that the same latent state would not be reselected.

#### Dynamics outside the PFC

Within-layer recurrent connectivity in the striatum, thalamus, and motor output is set to produce bump attractor dynamics, with local excitation and surround inhibition, following classical continuous attractor models (Ben-Yishai et al., 1995; Zhang, 1996). Each recurrent weight matrix *W* ∈ ℝ^*M ×M*^ is constructed by centering a Gaussian profile of width *w*_*b*_ on each neuron, normalizing total recurrent input from the excitatory region to unity, and setting inhibitory weights so that one active bump produces unit inhibition elsewhere.

This choice maximizes re-use of a common architecture, and is consistent with the topographic organization of these areas (Alexander et al., 1986; Frank & Claus, 2006; Gurney et al., 2001), but we wish to be clear that our treatment of the basal ganglia is an extreme simplification, and thus wrong in detail. For example, here we have used local recurrence as a means to create topography whereas in the brain we assume that the topography is inherited from inputs, and dependent on aspects of basal ganglia architecture omitted here for simplicity.

#### Projections between layers

The basal ganglia receives input:

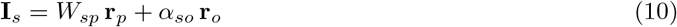

Here, *W*_*sp*_ is the learned weight matrix from the PFC and *α*_*so*_ scales a topographic motor output projection.

The basal ganglia projects to the thalamus, which receives:

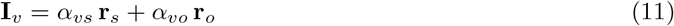

where *α*_*vs*_ and *α*_*vo*_ are scalar gains on the striatal and motor output projections. Both projections are topographically organized.

The motor output layer receives input from the thalamus and external supervision:

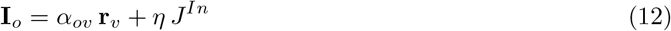

where *α*_*ov*_ is a scalar gain on the thalamic projection and *J*^*In*^ is the supervisory input vector, present only during the learning phase.

#### Plasticity

Synaptic plasticity in the SIAS model takes place within the PFC-to-striatum projection (*W*_*sp*_) and, in the flexible model, the feedback projections from motor cortex to PFC (*W*_*po*_). In both cases, we use the Hebbian learning rule noted in the text, with pre- and post-synaptic thresholds. Weights are updated during a learning period in each trial, are clipped to the range [*W*_min_, *W*_max_], and are subject to competitive dynamics that encourage topographic specificity.

More concretely, after each trial, competitive decay rules are applied. The changepoint model uses output competition only, whereas the flexible model uses input competition as well. Specifically, in the changepoint model, output competition limits the development of one-to-many mappings from attractors to actions, ensuring that, for example, one PFC latent state can track a drifting action correctly. In the flexible model, this same competition is applied, along with an analogous form of input competition, which helps retain one-to-one (as opposed to many-to-one PFC to BG connectivity). Without input competition, drift in attractor-to-action mappings would occasionally cause overlap in latent state coding for actions that are close in space, leading to an inability to revert to exactly one attractor on reversals. This situation is avoided the majority of the time without competition, but competitive weights fully prevent occasional edge cases.

We formulate output competition as follows: for each PFC neuron *i*, the peak target *j*^*∗*^ = arg max_*j*_ | (*W*_*sp*_)_*ij*_| is identified, and weights outside a window of 2*w*_*s*_ around *j*^*∗*^ are decayed:

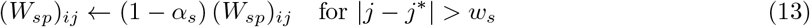

Individual neuron projections therefore stay localized in output space to approximately the standard prediction width.

Input competition, which applies to the flexible model only, is analogous: for each output neuron *j*, the dominant PFC group 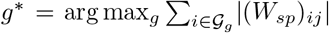 is identified, and all non-dominant groups’ weights to that neuron are decayed:

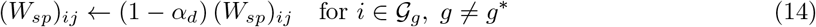

The flexible model additionally uses active-group decay on the plastic weights associated with the currently active PFC attractor. This mechanism limits the gradual spread of learned feedback weights and improves retrieval during reversals.

#### Within-trial prediction, supervision, and learning phases

Each trial proceeds through a prediction, supervision, and learning phase. In the decision phase, the network runs for *T*_dec_ steps without supervision, and the action is read out as the index of the maximally active motor output neuron at the end of the phase. In the supervision phase, *J*^*In*^ is applied to the motor output and the network runs for *T*_delay_ additional steps, allowing the supervised signal to propagate through the network before plasticity commences. A learning phase then runs for *T*_learn_ steps, during which time Hebbian learning is active.

Note that the synaptic learning rate *β* in Eq. (1) governs per-timestep weight increments during this phase; it is distinct from the trial-by-trial “learning rate” plotted in Figures 3–5, which reflects the behavioral ratio of prediction update to prediction error.

#### Supervision, thalamic surprise, and the adaptive threshold

The supervisory input *J*^*In*^ is a Gaussian-shaped vector of width *σ*_sup_ centered at the observed outcome location, scaled to a peak amplitude of *η* and offset by a uniform inhibitory baseline. This provides spatially localized excitation at the feedback location while suppressing activity elsewhere. This supervision propagates backwards to the thalamus, where it interacts with the feed-forward input to produce a surprise signal that controls the gating of PFC attractor states.

The surprise signal *S* computed over thalamic activities controls the PFC gating signal *δ* (Eqs. (2)– (3)). This signal measures the spatial spread of thalamic activity: when the proposed action and supervised feedback agree, thalamic activity is concentrated and *S* is low; when they conflict, activity is dispersed and *S* is elevated. When *δ* = 1, the gating signal suppresses the current PFC attractor via inhibition (Eq. (4)), facilitating a noise-driven transition to a new state.

Setting a single surprise threshold *S*_0_ for all subjects and conditions, however, does not allow the model to cope with varying noise levels. Therefore, we also implemented an adaptive threshold mechanism described in Eq. (5). As noted in the text, a running mean 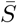 is updated after each trial using an exponential moving average with time constant *τ*_ema_.

### Helicopter task and performance analysis

We analyze human behavior and model performance using a cohort of 32 participants from a previously described predictive inference task (McGuire et al., 2014). In the task, an animated helicopter, occluded by clouds, drops bags of coins in an open field. Participants were pre-trained to move a bucket horizontally to catch falling bags. During the experiment, the helicopter occasionally changed (latent) locations abruptly at changepoints. Bag observations were sampled from Gaussian distributions with a standard deviation of 10 or 25, around the latent helicopter locations. The objective of the task was to catch as many bags as possible.

We fit the SIAS changepoint model to each of the 32 participants individually, optimizing a single free parameter, surprise threshold, over the range [0.06, 0.3] by minimizing the MSE between model and subject peri-changepoint learning rates (Figure 3A). Learning rates were computed as a function of post-changepoint trial position (positions 0–4 after each changepoint) by regressing trial-by-trial updates against prediction errors at each position, pooled across all changepoints for each participant.

We fit linear models, following earlier work on changepoint tasks (Nassar et al., 2010, 2012), to both subject and network-based behavior. Let *u*_*t*_ denote the trial-by-trial update, *p*_*t*_ the signed prediction error, *c*_*t*_ the changepoint probability (CPP), and *r*_*t*_ the relative uncertainty (RU). CPP and RU were centered before forming interaction terms, such that 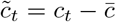 and 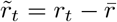. We fit the regression

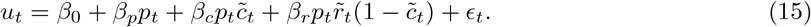

Here, *β*_*p*_ captures baseline prediction-error-driven updating, whereas *β*_*c*_ and *β*_*r*_ capture modulation of prediction-error-driven updating by CPP and RU, respectively. This form follows the logic of prior changepoint analyses in which CPP and RU influence learning by scaling the impact of prediction errors, while matching the centered-predictor implementation used in the present analyses.

Changepoint probability and relative uncertainty themselves are statistics derived from normative models of our tasks (Figure 5). In a highly accurate approximation of the full Bayesian model, updates depend on these terms linearly (Figure 3C). Intuitively, CPP reflects the probability of a change in helicopter location given the most recent observation, and RU measures the uncertainty about the helicopter’s location, assuming the next trial is not a changepoint. The normative model takes additional environment information (i.e., hazard rate and noise level) and these two latent variables to capture human learning behavior. More information can be found in the original papers (Nassar et al., 2010, 2012).

When fitting the linear model to human data, we extract prediction errors and updates, compute CPP and RU from the environmental parameters (hazard rate and noise level), and fit equation (15) using ordinary least square with centered CPP and RU regressors. For the model data, we run the SIAS model at each participant’s best-fit threshold on the same task and fit the same linear model to the resulting behavior (Figure 3C).

### Adaptive threshold fitting

For the SIAS model with an adaptive threshold, the surprise threshold follows equation (5). To characterize noise adaptation (Figure 5B), we ran the model on short tasks consisting of eight stable trials followed by two changepoint trials, across noise levels *σ*∈ {5, 10, 25} and prediction errors ranging from 0 to 100. We computed the learning rate on the final trial as the ratio of update magnitude to prediction error magnitude, averaged over 50 repetitions per condition. The reduced Bayesian model was evaluated in the same manner (Figure 5A).

We then applied the adaptive-threshold SIAS model to human data (Figure 5C). Learning rates were computed by binning absolute prediction errors and averaging the corresponding update-to-error ratios within each bin, separately for each noise level (*σ* ∈ *{*10, 25*}*, matching the experimental conditions).

### Applying the model to mixed changepoint and reversal environments

To evaluate performance in mixed environments, we generated 100-trial tasks with context changes every 10 trials, yielding 10 contexts. Latent states were generated monotonically with a minimum separation of 55 units, and observations were sampled from Gaussian distributions with *σ* = 5 centered on each latent state. The first two contexts were always novel changepoints; the remaining eight context changes contained a controlled number of reversals (*n*_rev_ ∈ {0, 1, …,8}), selected uniformly without replacement from the eligible post-warmup context changes. On a reversal, the latent state reverted to a uniformly selected previously visited non-current state.

We ran 1000 repetitions per condition for both the changepoint and flexible models and computed post-transition absolute prediction error relative to the latent distribution mean, excluding the first two warm-up contexts. The number of inferred states (Figure 6C) was measured by counting distinct state-action associations in the PFC-to-striatum weight matrix at the end of each simulation.

## Acknowledgements

This work was supported by National Center for Artificial Intelligence CENIA FB210017, Basal ANID and Daniel Scott was supported by NIMH training grant T32MH115895.

